## Supplement figures for "The CD4 transmembrane GGXXG and juxtamembrane (C/F)CV+C motifs mediate pMHCII-specific signaling independently of CD4-LCK interactions"

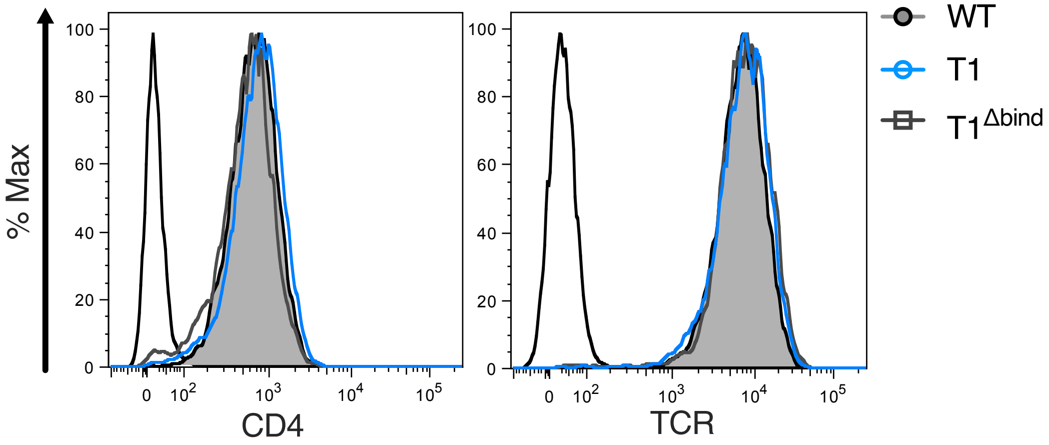


**Figure 2 — figure supplement 1**

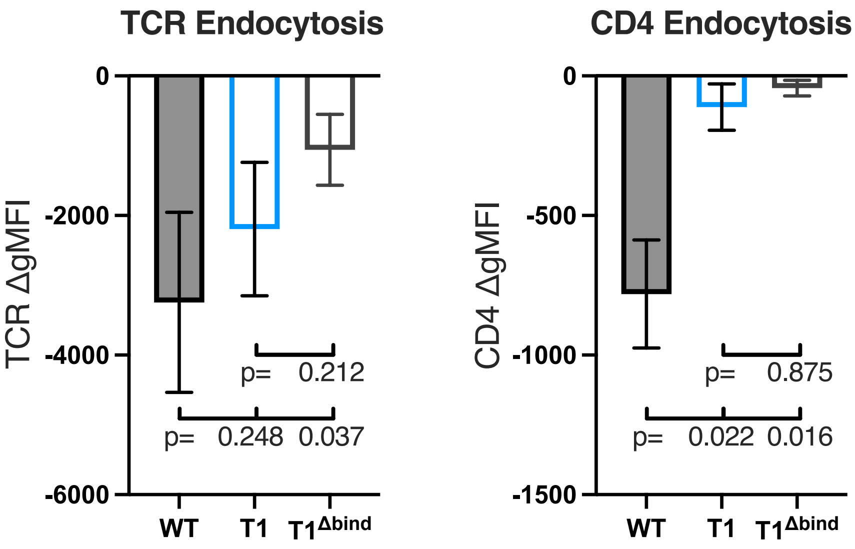


Figure 2 — figure supplement 2

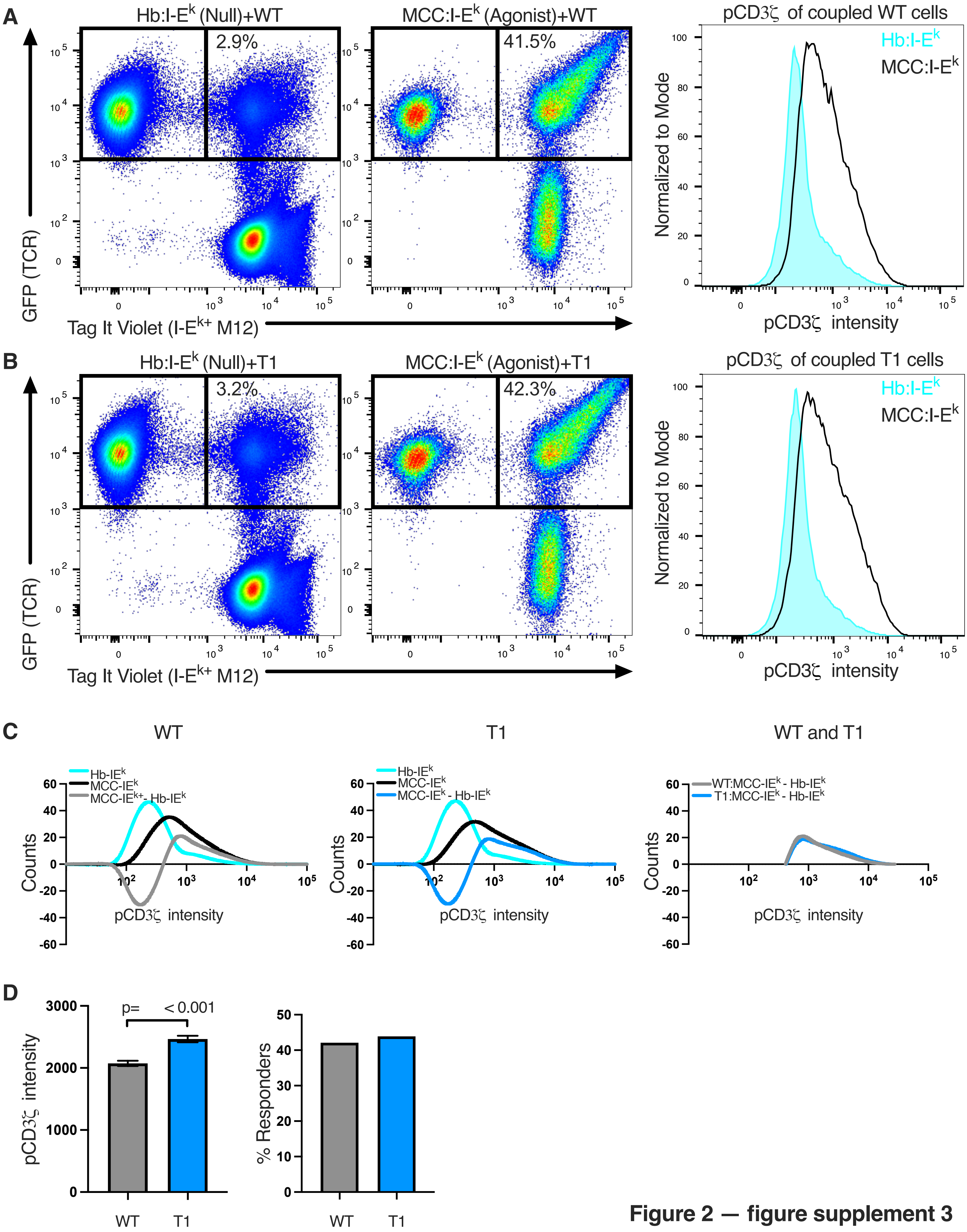


**Figure 2 — figure supplement 3**

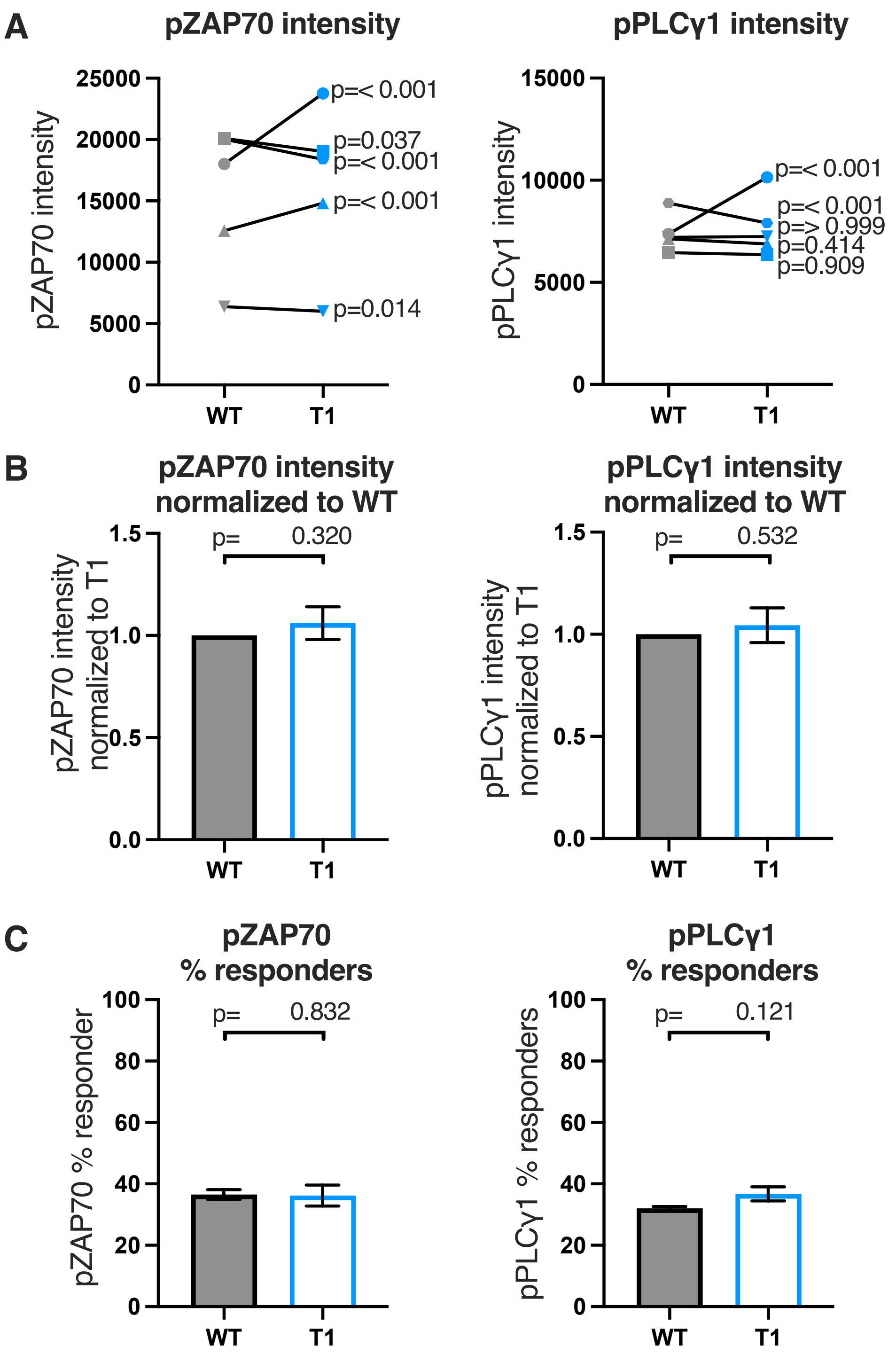


**Figure 2 — figure supplement 4**

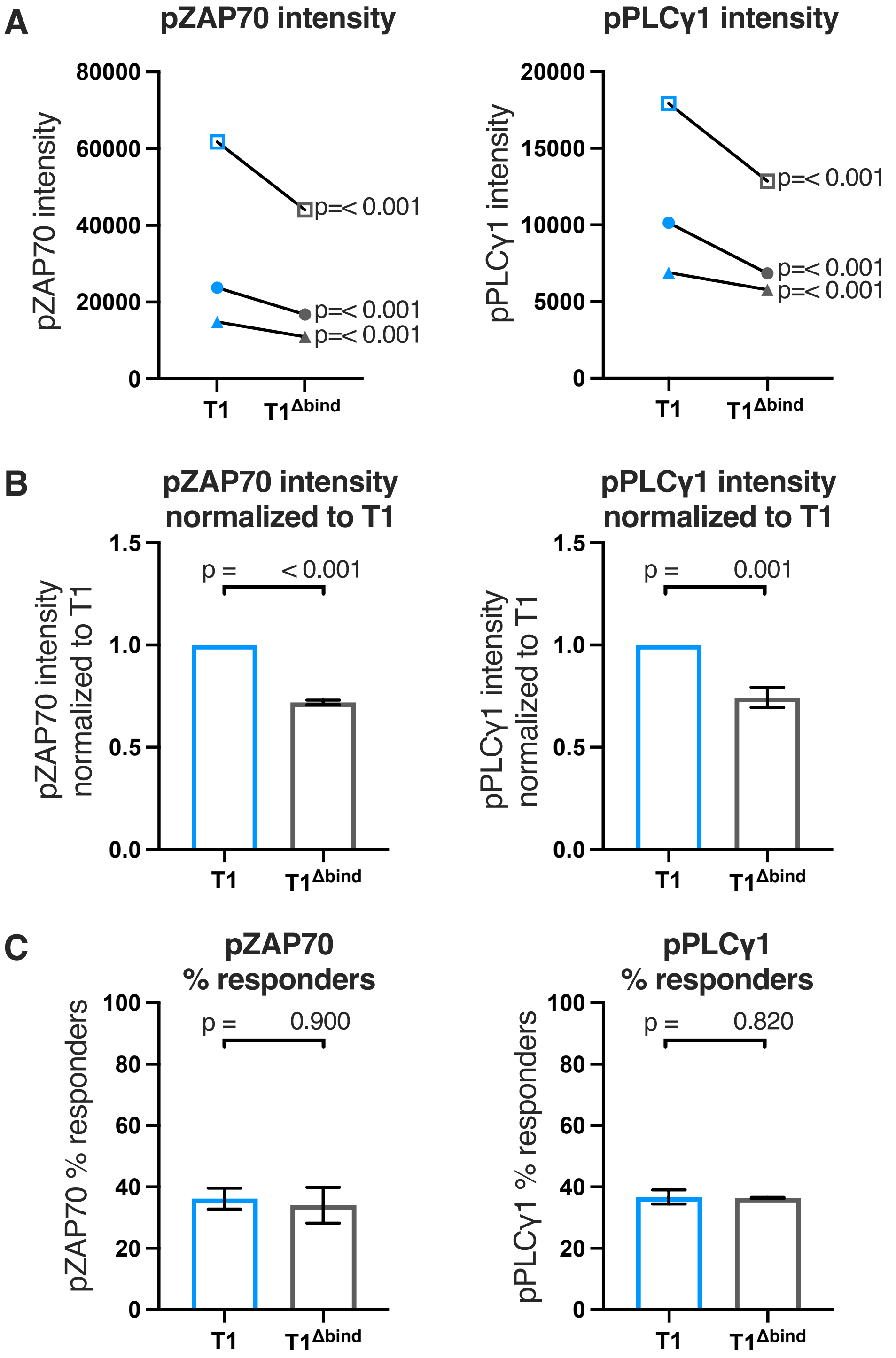


**Figure 2 — figure supplement 5**

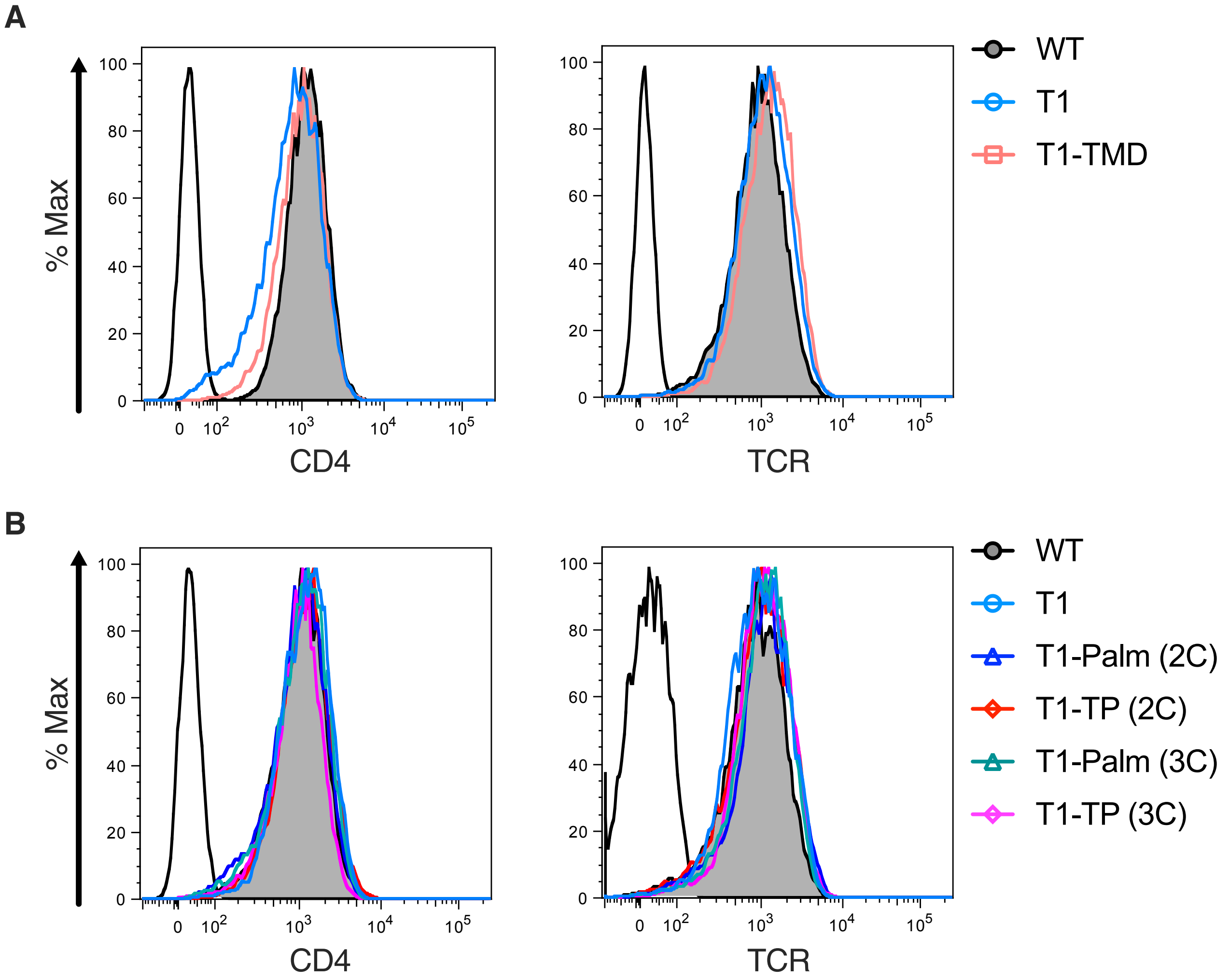


**Figure 3 — figure supplement 1**

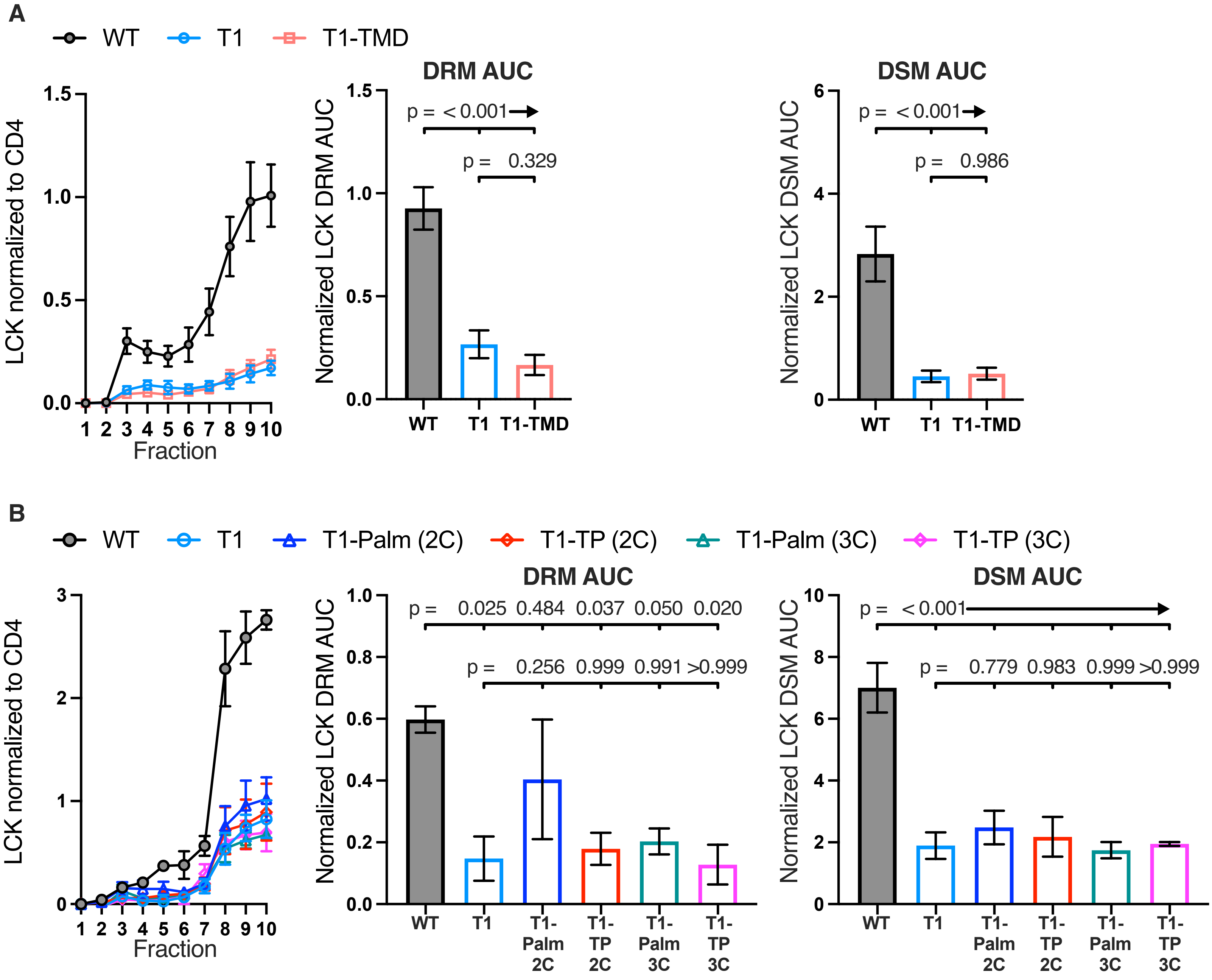


Figure 3 — figure supplement 2

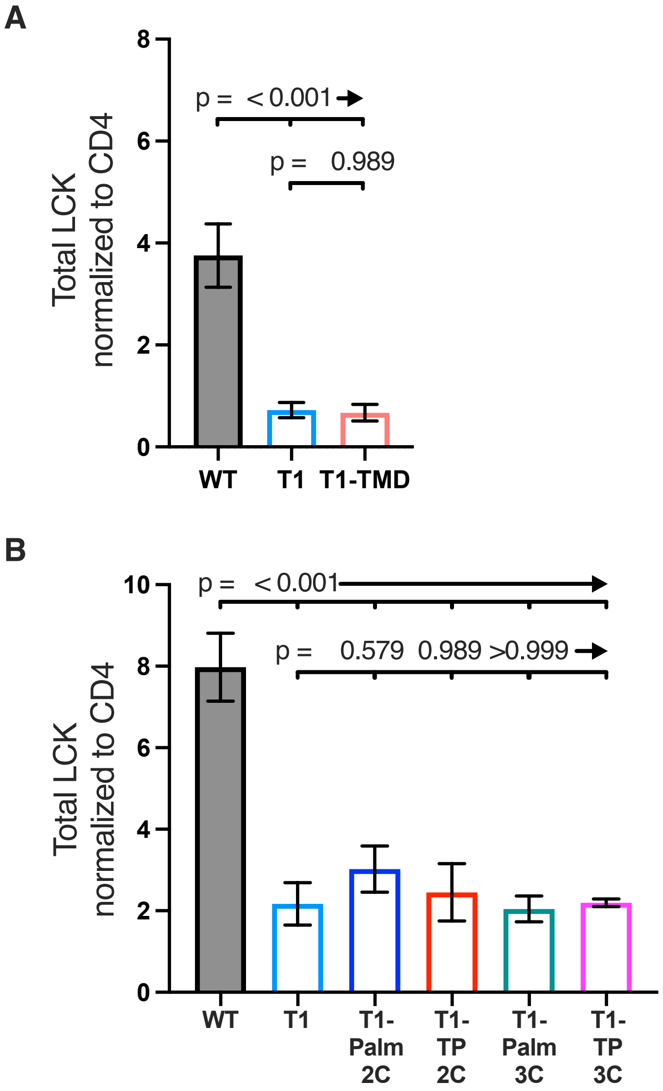


**Figure 3 — figure supplement 3**

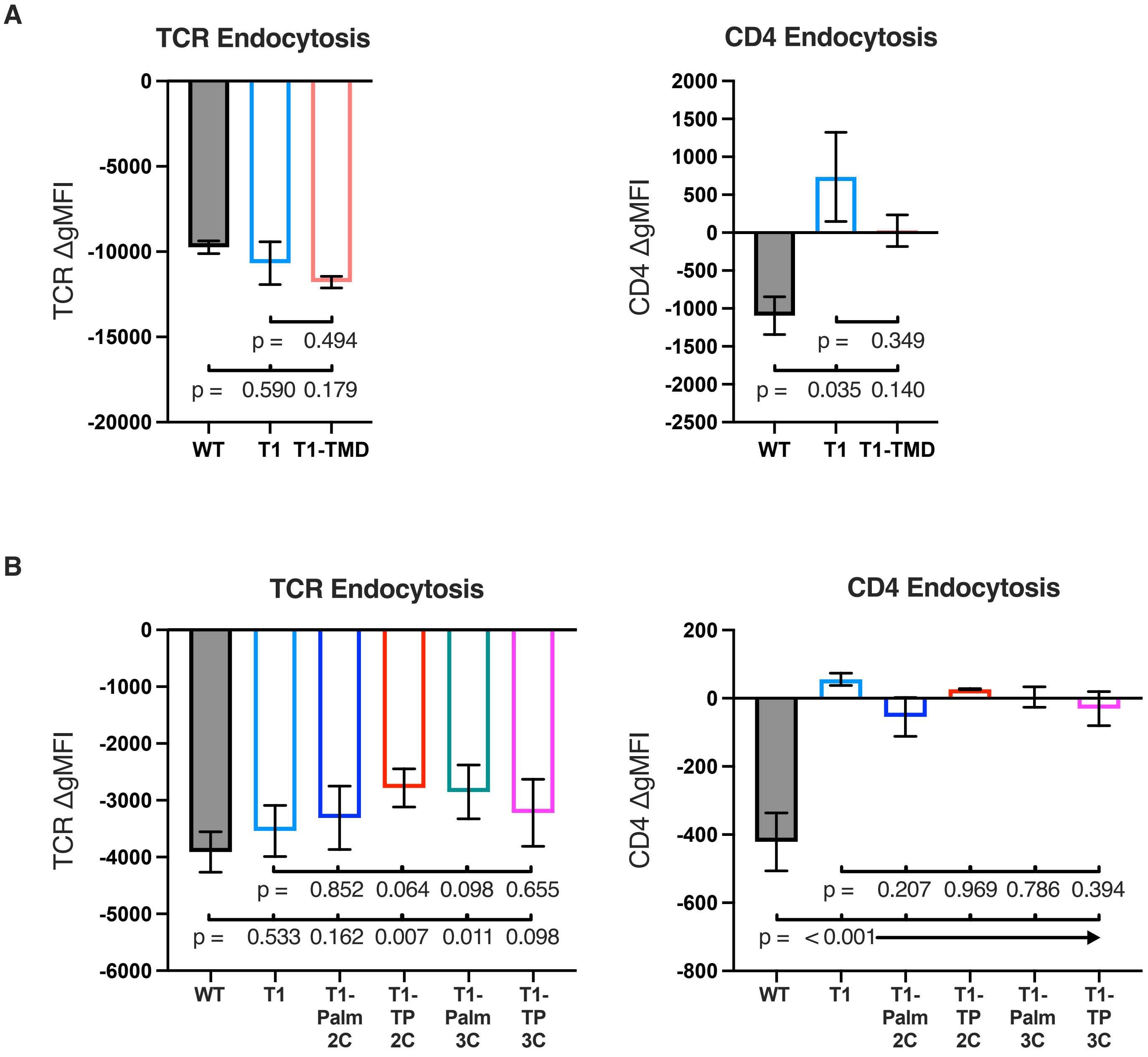


**Figure 3 — figure supplement 4**

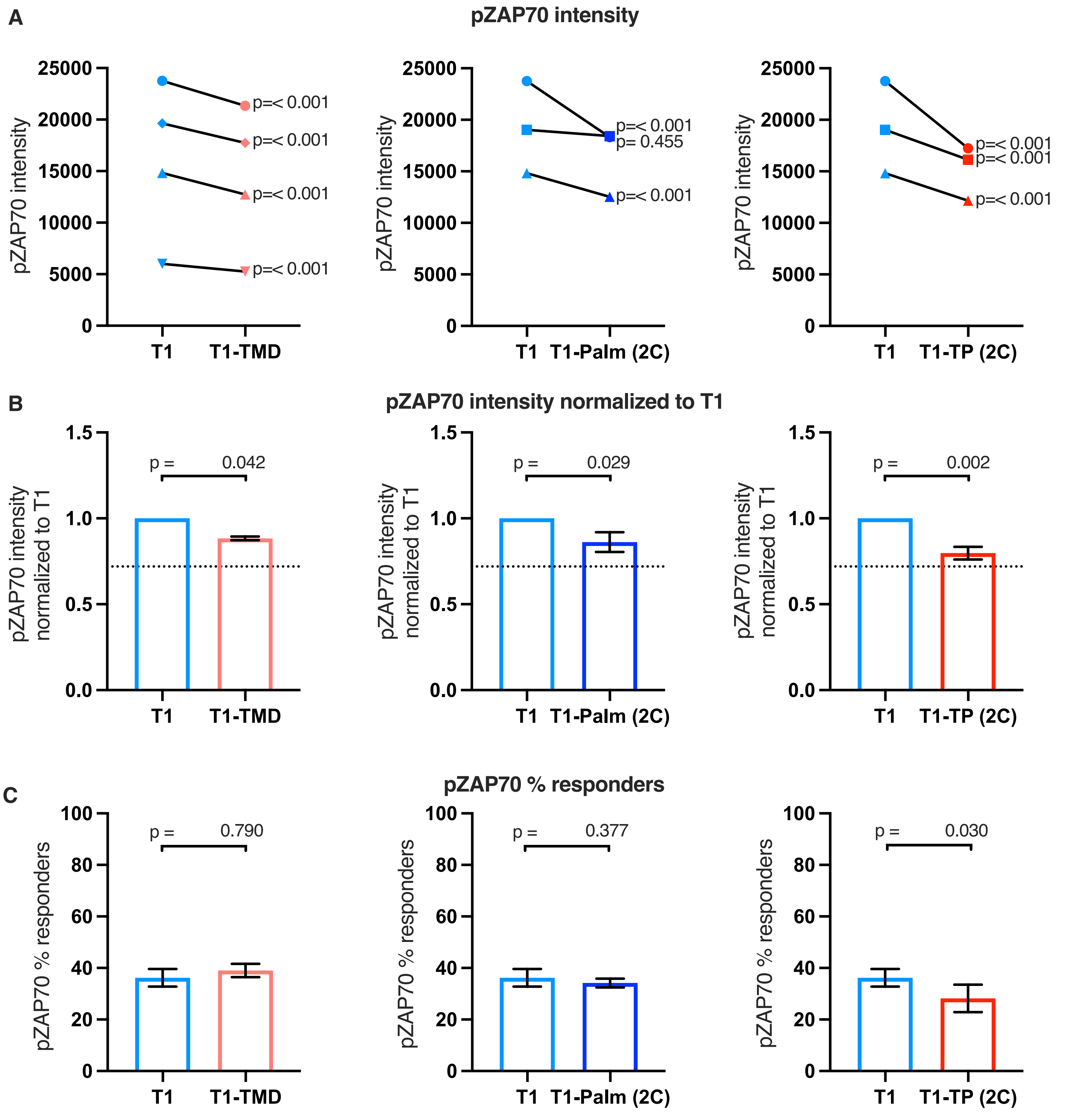


Figure 4 — figure supplement 1

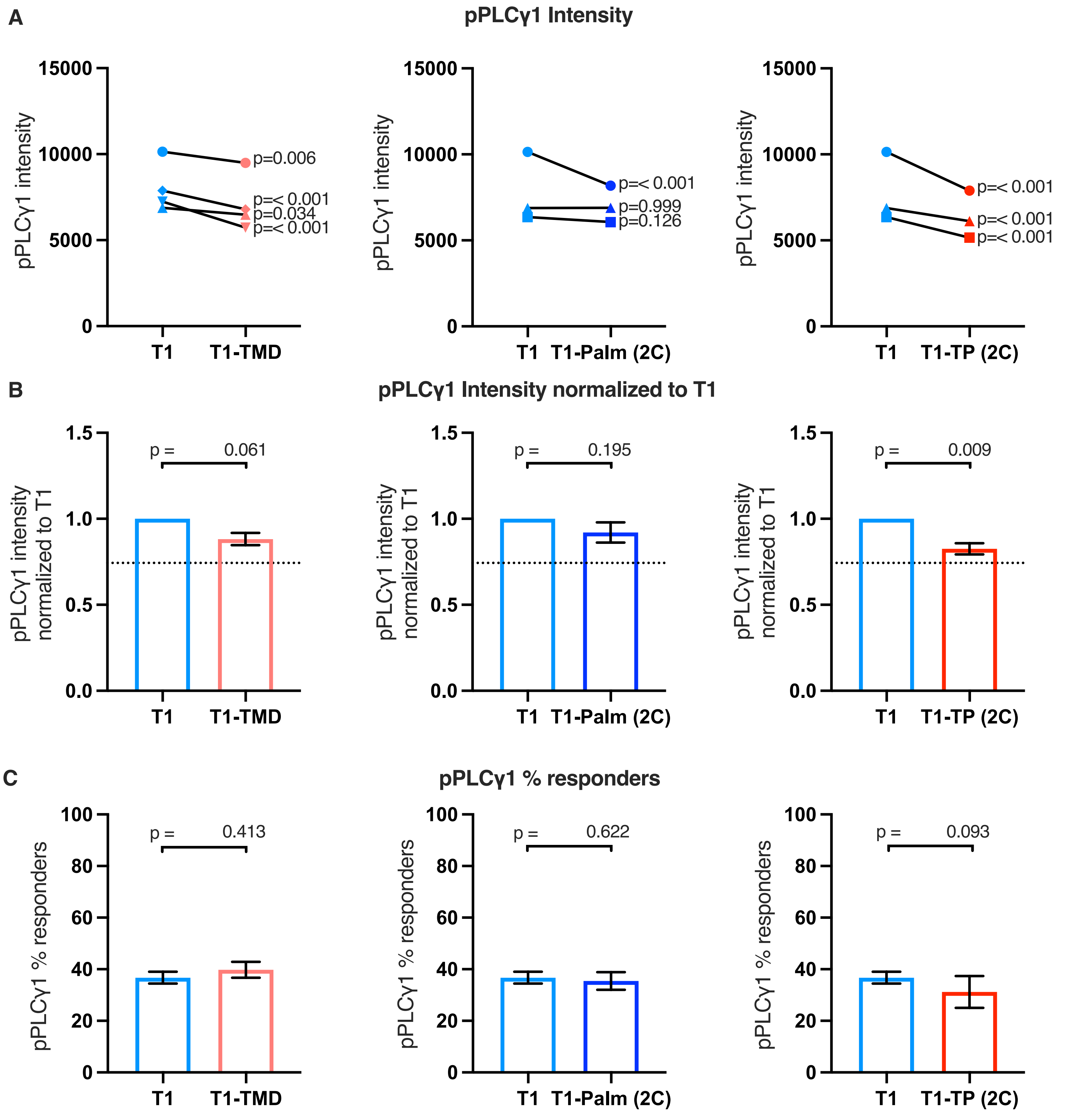


Figure 4 — figure supplement 2

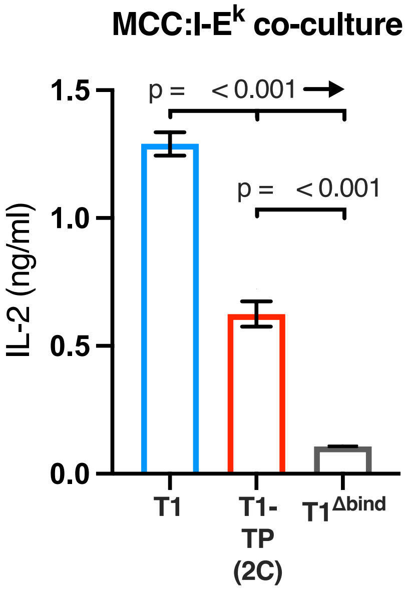


**Figure 4 — figure supplement 3**

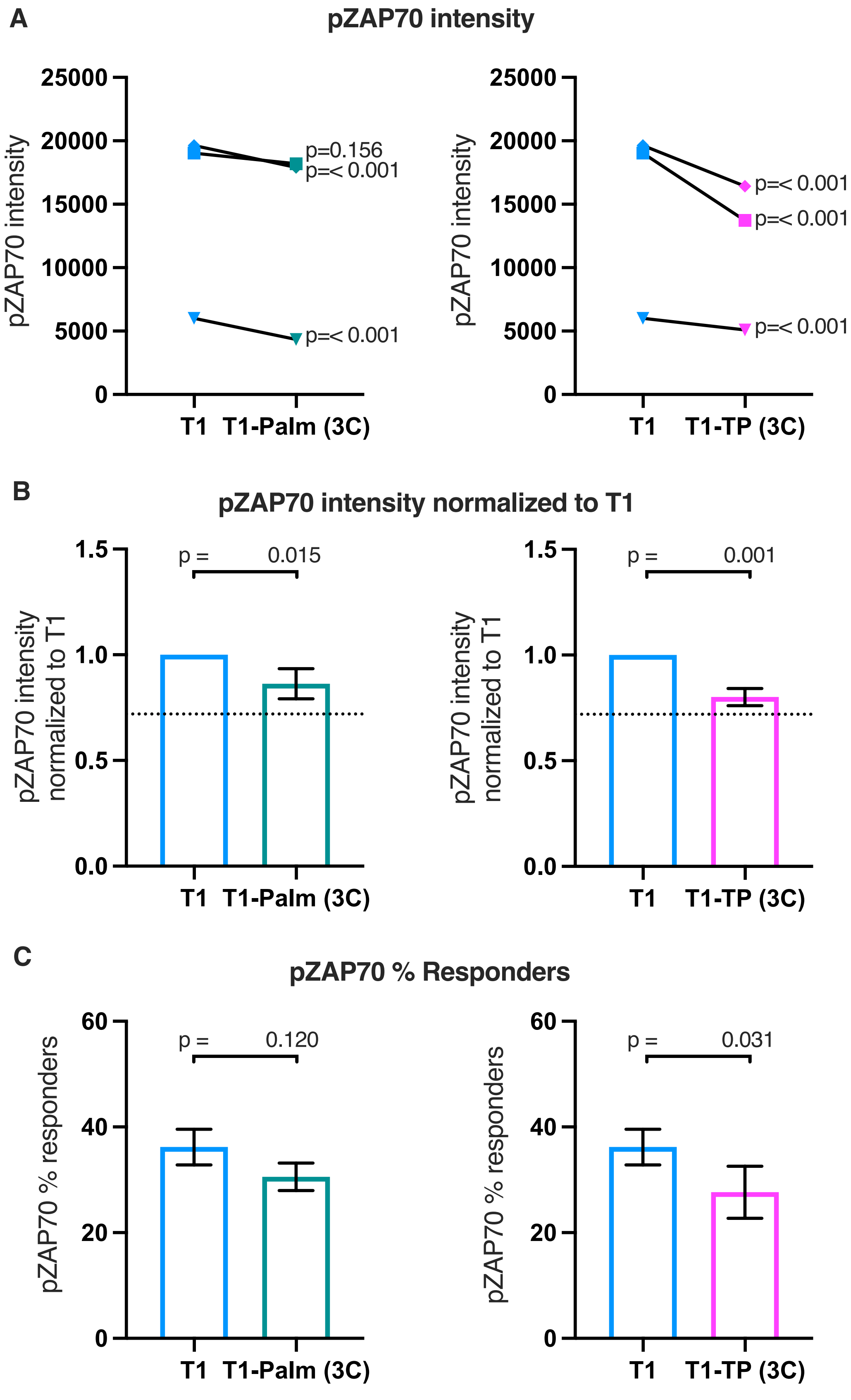


Figure 5 — figure supplement 1

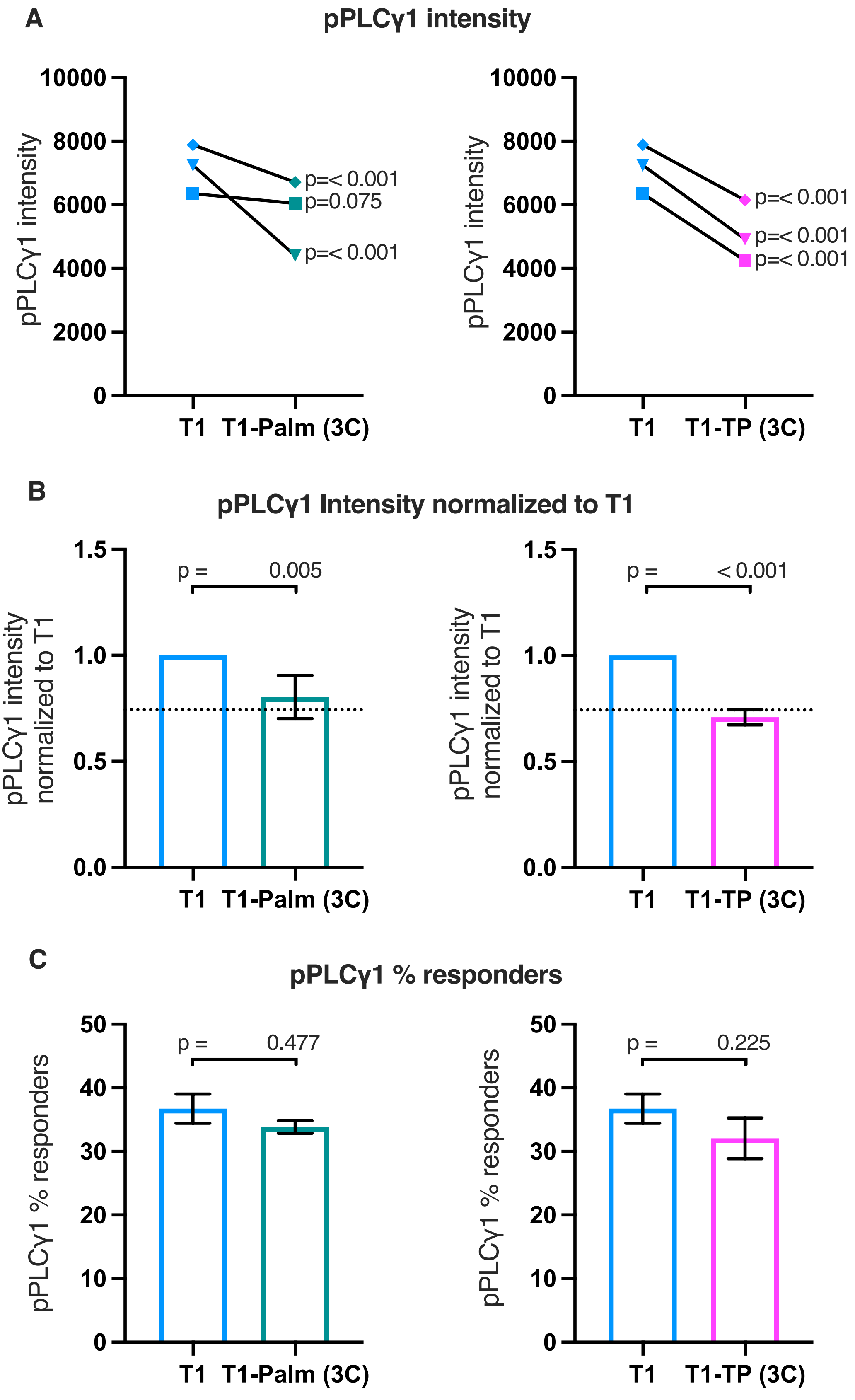


Figure 5 — figure supplement 2

### Figure Supplement Legends

**Figure 2- figure supplement 1.**
Flow cytometry analysis of CD4 (left) and TCR (right) expression on $58\alpha^-\beta^-$ hybridoma cells. Parental $58\alpha^-\beta^-$ hybridoma cells served as negative control for surface expression (open black histogram trace).

**Figure 2- figure supplement 2.**
TCR (left) and CD4 (right) endocytosis after pMHCII engagement is shown for the indicated cell lines after 16 hours coculture with APCs in the presence of $10\mu M$ MCC peptide. The change in TCR and CD4 gMFI, as measured by flow cytometry, is shown for each cell line relative to an equivalent sample cultured with APCs in the absence of MCC peptide. Each data point represents the mean ± SEM for three independent experiments. For endocytosis, measurements were performed in triplicate for each experiment. One-way ANOVA was performed with a Dunnett's posttest.

**Figure 2-figure supplement 3.**
Example of intracellular signaling analysis workflow.
(**A**) Flow cytometry analysis of WT $58\alpha^-\beta^-$ hybridoma:M12 cell couples. Representative dot plots are shown for $TCR\beta^{GFP+}$ $CD4^+$ $58\alpha^-\beta^-$ hybridoma cells coupled to Tag-it Violet-labeled M12s expressing the indicated tethered null Hb:I-$E^k$ (left) and agonist MCC:I-$E^k$ pMCHII (center). Representative histograms of WT $58\alpha^-\beta^-$ hybridoma cells coupled to M12 cells transduced to express the indicated tethered pMHCII are shown for pCD3$\zeta$ intensity (right). 10,000 coupled cells were collected per individual experiment.
(**B**) Flow cytometry analysis of T1 $58\alpha^-\beta^-$ hybridoma-M12 cell couples. Representative dot plots are shown for $TCR\beta^{GFP+}$ $CD4^+$ $58\alpha^-\beta^-$ hybridoma cells coupled to Tag-it Violet-labeled Hb:I-$E^{k+}$ (left) and MCC:I-$E^{k+}$ (center) M12 cells. Representative histograms of T1 $58\alpha^-\beta^-$ hybridoma cells coupled to M12 cells expressing the indicated tethered pMHCII are shown for pCD3$\zeta$ intensity (right). 10,000 coupled cells were collected per individual experiment.
(**C**) A representative smoothed overlapping histogram of pCD3$\zeta$ intensity is shown for $58\alpha^-\beta^-$ cells coupled to Hb:I-$E^{k+}$ (cyan) or MCC:I-$E^{k+}$ (black) M12 cells. Histogram of pCD3$\zeta$ intensity for $58\alpha^-\beta^-$ cells coupled to MCC:I-$E^{k+}$ M12 cells subtracted from Hb:I-$E^{k+}$ M12 cell couples show the difference in pCD3$\zeta$ intensity on a bin-by-bin basis after stimulation with agonist MCC:I-$E^k$ compared with null Hb:I-$E^k$ for WT (left) and T1 (center) cells. Overlapping pCD3$\zeta$ histogram (right) of cells responding to MCC:I-$E^k$ after Hb:I-$E^k$ subtraction shows the responding populations for the WT and T1 cell lines. Data represent the aggregate from three individual experiments (30,000 couples analyzed). One-way ANOVA was performed with a Dunnett's posttest for comparison with the WT sample because other mutants were simultaneously analyzed in this experiment (not shown).
(**D**) Concatenated pCD3$\zeta$ average intensity ± SEM of WT and T1 cells (left) and the percent of responding WT and T1 cells (right).

**Figure 2-figure supplement 4.**

(**A**) ZAP70 (left), and PLC$\gamma$1 (right) phosphorylation intensity for WT and T1 are shown for paired (connecting line) WT and T1 cell lines. Five independently generated cell lines were tested (biological replicates). For phosphorylation intensity, each pair of lines (connected symbols) was tested in three independent experiments (technical replicates). The data from those experiments was aggregated, and the symbols represent the mean intensity of the aggregated pCD3$\zeta$ intensity values. One-way ANOVA was performed with a Dunnett's posttest.
(**B**) Average T1 phosphorylation for ZAP70 (left) and PLC$\gamma$1 (right) from A are shown normalized to the paired WT controls. Bars represent the mean +/- SEM. One-way ANOVA was performed with a Sidak's posttest for specific comparisons.
(**C**) Average WT and T1 % responders for pZAP70 (left), and pPLC$\gamma$1 (right) are shown for the cell lines shown in A. Bars represent the mean +/- SEM. One-way ANOVA was performed with a Sidak's posttest for specific comparisons.

**Figure 2-figure supplement 5.**
(**A**) Phosphorylation intensity for T1 and T1$^{\Delta bind}$ ZAP70 (left), and PLC$\gamma$1 (right) are shown for paired (connecting line) T1 and T1$^{\Delta bind}$ cell lines. Three independently generated cell lines were tested (biological replicates). Analysis was performed as in Figure 2— figure supplement 4 with the exception that the open symbol represent data from a single experiment whereas the closed symbols represent the average of aggregated data from three independent experiments. For the open symbol comparisons we performed a two-tailed t test as no other samples were collected in parallel.
(**B**) Average T1$^{\Delta bind}$ cell line phosphorylation intensity for ZAP70 (left), and PLC$\gamma$1 (right) are shown normalized to the average intensity of the paired T1 control cells shown in A. Analysis was performed as in Figure 2— figure supplement 4.
(**C**) Average T1 and T1$^{\Delta bind}$ cell line % responders for pZAP70 (left) and pPLC$\gamma$1 (right) are shown. Analysis was performed as in Figure 2— figure supplement 4.

**Figure 3- figure supplement 1.**
(**A, B**) Flow cytometry analysis of CD4 (left) and TCR (right) expression on 58$\alpha^-\beta^-$ hybridoma cells. Parental 58$\alpha^-\beta^-$ hybridoma cells served as negative control for surface expression (open black histogram trace).

**Figure 3- figure supplement 2.**
(**A, B**) LCK signal is shown for each sucrose fraction normalized to the CD4 signal detected in the corresponding fraction (left). The AUC is shown for the normalized LCK signal in the DRM (center) and DSM (right) fractions. Each data point represents the mean +/- SEM for three independent experiments with the same cell line. Data are representative of experiments performed with three independently generated sets of lines for TMD, Palm(3C), and TP(3C) mutants. Analysis was performed with two set of lines for the Palm(2C) and TP(2C) mutants. One-way ANOVA was performed with a Dunnett's posttest for comparisons with WT and T1 samples, and a Sidak's posttest for comparisons between selected samples.

**Figure 3- figure supplement 3.**

(**A, B**) Total LCK signal (total AUC for sucrose gradient) normalized to CD4 signal is shown for the indicated cell lines. Each data point represents the mean +/- SEM for three independent experiments with the same cell line (experimental replicates). Data are representative of experiments performed with three independently generated sets of lines for TMD, Palm(3C), and TP(3C) mutants (biological replicates). Analysis was performed with two set of lines for the Palm(2C) and TP(2C) mutants. One-way ANOVA was performed with a Dunnett's posttest for comparisons with WT and T1 samples, and a Sidak's posttest for comparisons between selected samples.

**Figure 3- figure supplement 4.**
(**A, B**) TCR (left) and CD4 (right) endocytosis after pMHCII engagement is shown for the indicated cell lines after 16 hours coculture with APCs in the presence of 10μM MCC peptide. The change in TCR and CD4 gMFI, as measured by flow cytometry, is shown for each cell line relative to an equivalent sample cultured with APCs in the absence of MCC peptide. Each data point represents the mean ± SEM for three independent experiments (experimental replicates). Data are representative of those acquired with at least two independently generated sets of cell lines (biological replicates). Endocytosis measurements were performed in triplicate (technical replicates) for each experiment. One-way ANOVA was performed with a Dunnett's posttest for comparisons with WT and T1 samples.

**Figure 4- figure supplement 1.**
(**A**) Phosphorylation intensity of ZAP70 for T1 and T1-TMD (left), T1 and T1-Palm (2C) (center), and T1 and T1-TP (2C) (right) are shown for paired (connecting line) cell lines. Analysis was performed as in Figure 4A.
(**B**) Normalized phosphorylation intensity of ZAP70 for T1-TMD (left), T1-Palm (2C) (center), and T1-TP (2C) (right) are shown as bars. Analysis was performed as in Figure 4B.
(**C**) Average % responders of the phosphorylation ZAP70 for T1-TMD (left), T1-Palm (2C) (center), and T1-TP (2C) (right) are shown as bars compared to the average of their paired T1 cell lines. Analysis was performed as in Figure 4C.

**Figure 4- figure supplement 2.**
(**A**) Phosphorylation intensity of PLCγ1 for T1 and T1-TMD (left), T1 and T1-Palm (2C) (center), and T1 and T1-TP (2C) (right) are shown for paired (connecting line) cell lines. Analysis was performed as in Figure 4A.
(**B**) Normalized phosphorylation intensity of PLCγ1 for T1-TMD (left), T1-Palm (2C) (center), and T1-TP (2C) (right) are shown as bars. Analysis was performed as in Figure 4B.
(**C**) Average % responders of phosphorylated pPLCγ1 for T1-TMD (left), T1-Palm (2C) (center), and T1-TP (2C) (right) are shown as bars compared to the average of their paired T1 cell lines. Analysis was performed as in Figure 4C.

**Figure 4- figure supplement 3.** Representative IL-2 production is shown in response to M12 cells transduced to express tethered MCC:I-$E^k$ constructs as used in phosphorylation analysis. Experiments were performed in triplicate and each bar equals the mean +/- SEM at that peptide concentration. Results are representative of those obtained with two independently generated matched sets of cell lines. One-way ANOVA was performed with a Dunnett's posttest for comparisons with WT and T1 samples, and a Sidak's posttest for comparisons between selected samples.

**Figure 5- figure supplement 1.**
(**A**) Phosphorylation intensity of ZAP70 for T1 and T1-Palm (3C) (left) and T1 and T1-TP (3C) (right) are shown for paired (connecting line) cell lines. Analysis was performed as in Figure 5A.
(**B**) Normalized phosphorylation intensity of ZAP70 for T1-Palm (3C) (left) and T1-TP (3C) (right) are shown as bars. Analysis was performed as in Figure 5B.
(**C**) Average % responders of phosphorylated ZAP70 for T1-Palm (3C) (left) and T1-TP (3C) (right) are shown as bars compared to the average of their paired T1 controls. Analysis was performed as in Figure 5C.

**Figure 5- figure supplement 2.**
(**A**) Phosphorylation intensity of PLC$\gamma$1 for T1 and T1-Palm (3C) (left) and T1 and T1-TP (3C) (right) are shown for paired (connecting line) cell lines. Analysis was performed as in Figure 5A.
(**B**) Normalized phosphorylation intensity of PLC$\gamma$1 for T1-Palm (3C) (left) and T1-TP (3C) (right) are shown as bars. Analysis was performed as in Figure 5B.
(**C**) Average % responders of phosphorylated PLC$\gamma$1 for T1-Palm (3C) (left) and T1-TP (3C) (right) are shown as bars compared to the average of their paired T1 controls. Analysis was performed as in Figure 5C.
